## Supplementary Information for "Deconstructing body axis morphogenesis in zebrafish embryos using robot-assisted tissue micromanipulation"

### Supplementary Figures

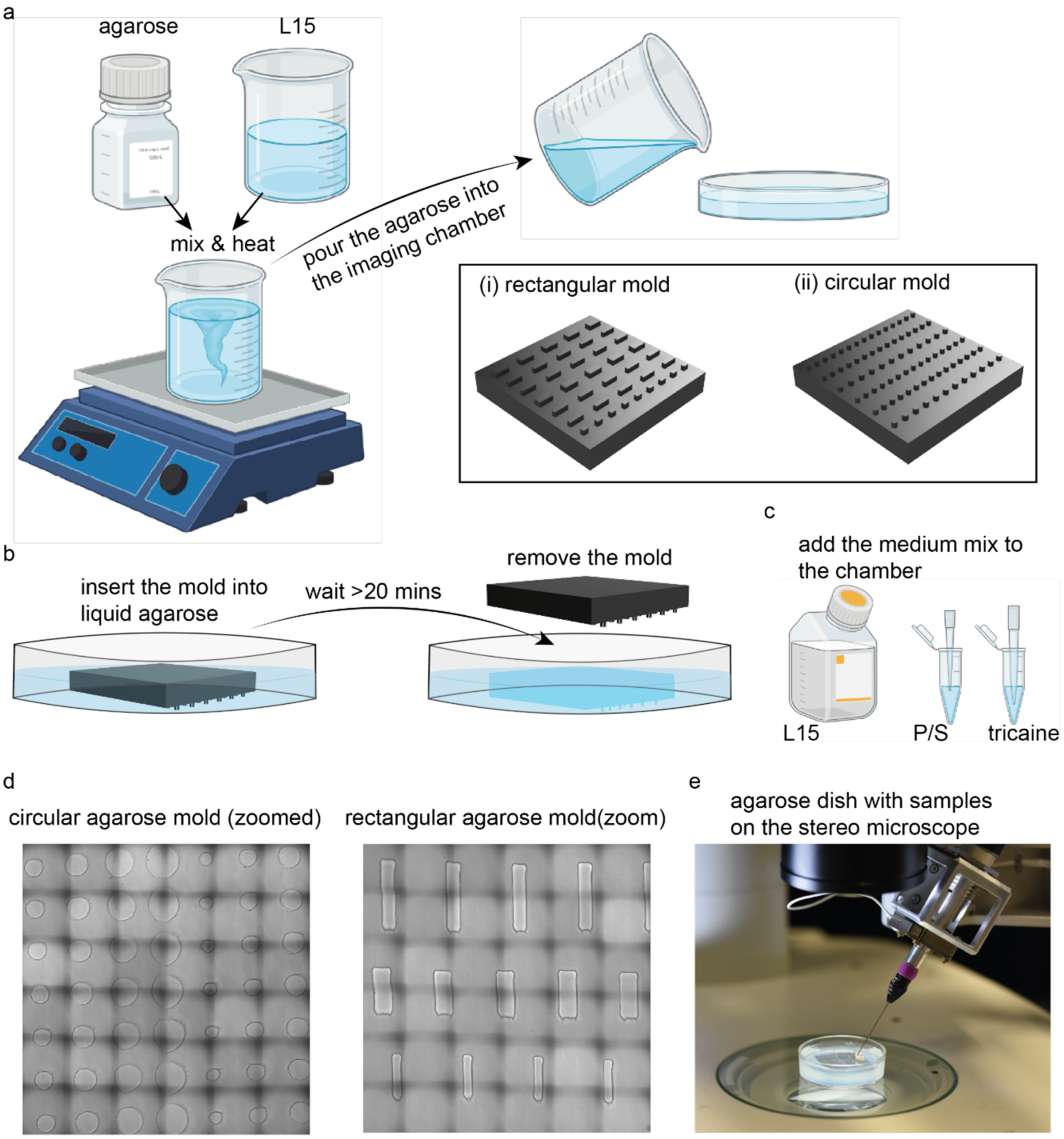


**Supplementary Fig. 1 Chamber preparation process for the robotic micromanipulations on the stereoscope and imaging on Nikon Ti microscope. a** Preparation of the agarose gel. **b** Preparation of the micromanipulation and/or imaging chambers. **c** Imaging medium components. **d** Bright field image of the microengineered manipulation and imaging chambers. **e** Manipulation chamber on the stereomicroscope for microsurgery.

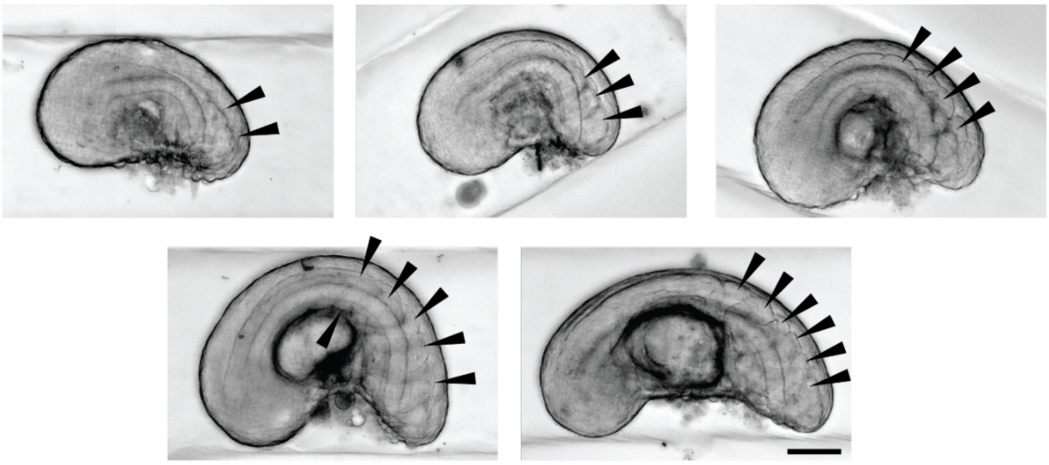

**Supplementary Fig. 2 Explants with 2 to 6 somites on them right after the robotic microsurgery.** Black arrows on the BF images (Gaussian based stack focuser in Fiji is used) of the tail explants indicate the individual somites on the tail explants which are acquired with robotic microsurgery. Scale bar, 100 μm.

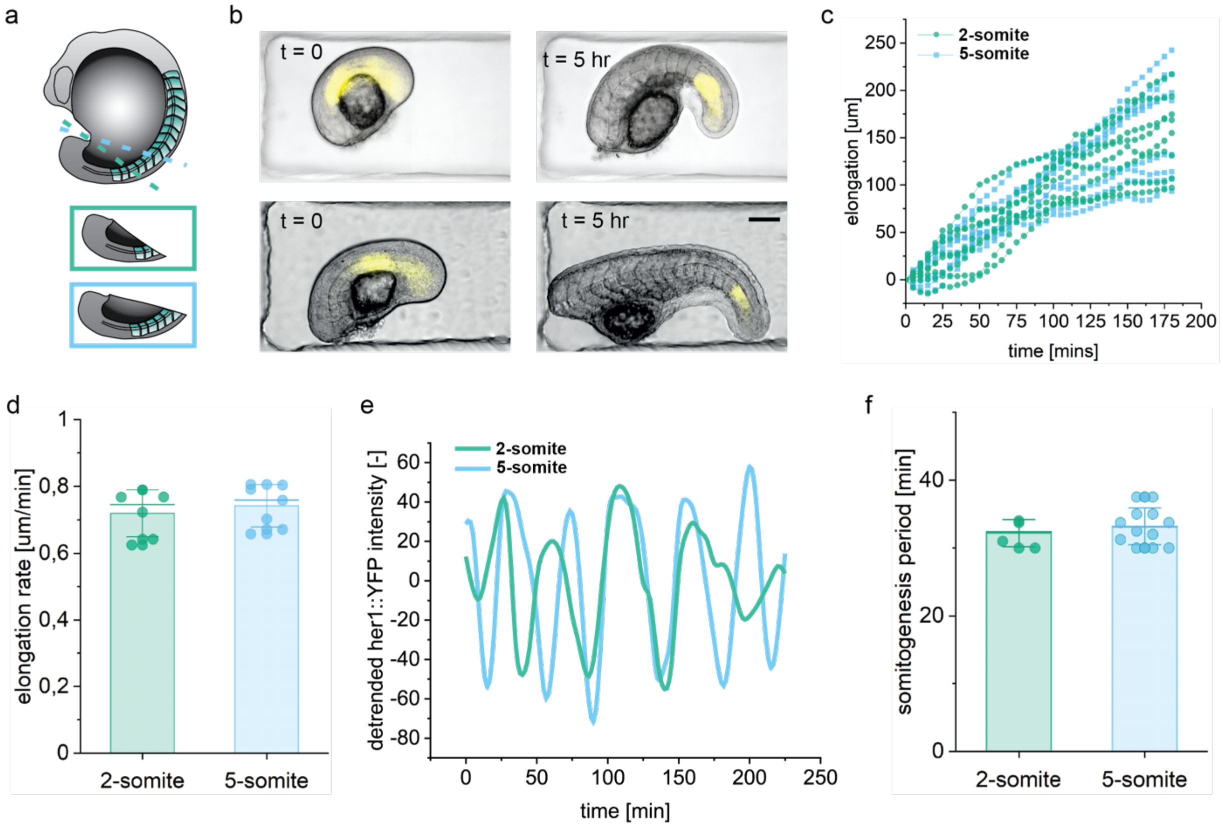


**Supplementary Fig. 3 Comparison of explants containing different number of somites. a** Schematic showing 2 and 5 somite explants. **b** Composite BF and YFP images for t = 0 and 5 hours of 2-somite explant (top) and 5-somite explant (bottom). Scale bar, 100 μm. **c** Elongation profile of several samples of 2-somite (n = 6) and 5-somite explants (n = 7). **d** Scatter point plot of elongation rate for 2-somite (n=6) and 5-somite explants (n=7). **e** Example curve of Her1-YFP intensity at the anterior for 2 and 5-somite explants. **f** Scatter point plot of anterior period of 2-somite (n=4) and 5-somite explants (n=12).

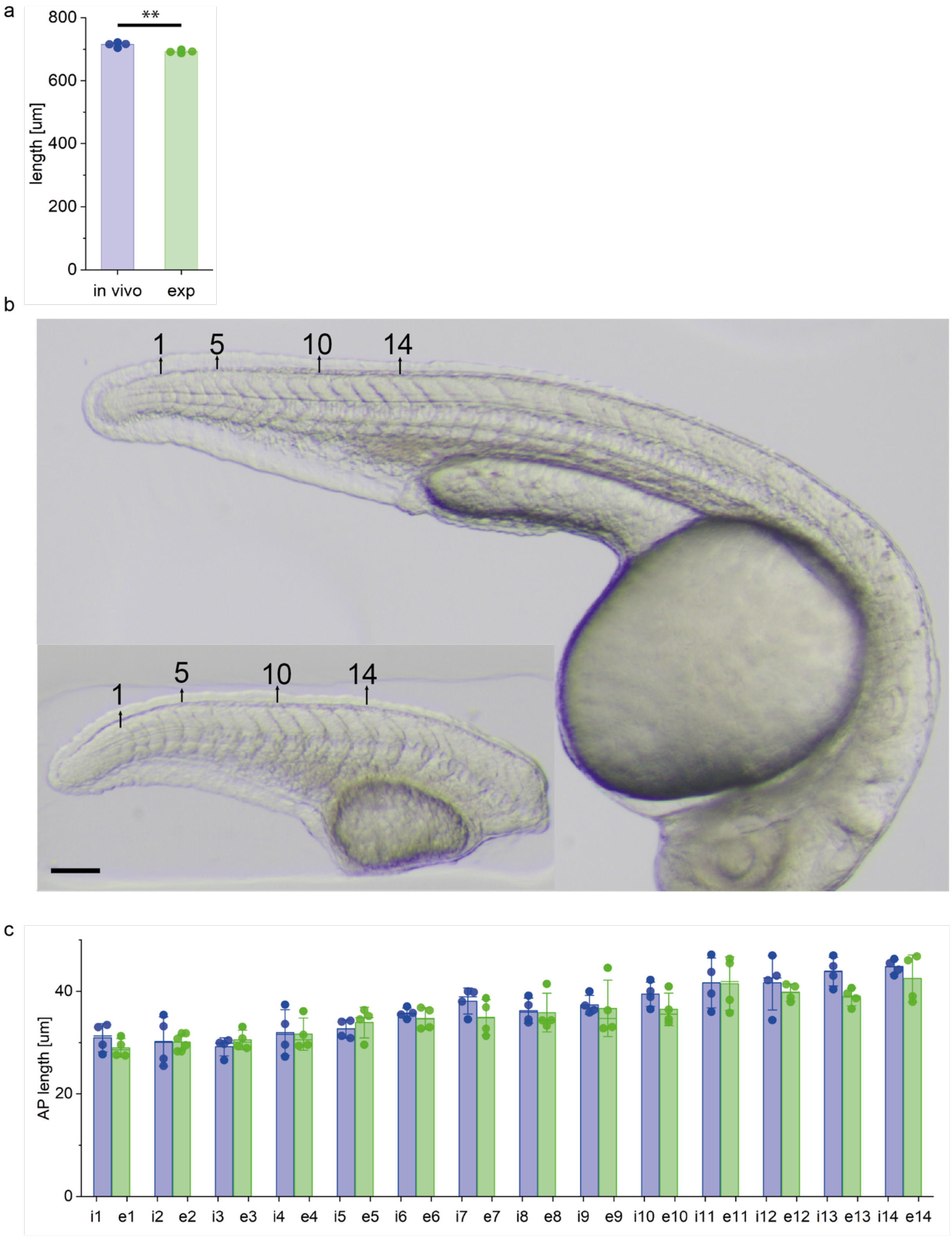

**Supplementary Fig. 4 Somite size comparison of in vivo and explant. a** Final total length of the *in vivo* and explant (n = 4, 4). Measurements are taken right after the end of the somitogenesis of the *in vivo* samples. **b** Example image of an i*n vivo* and explant to show the stage of the samples during the measurements indicating the numbering of the somites. **c** Somite size measurements of the last formed 14 somites for in vivo and explant (n = 4, 4). “i” and “e” in the x-axis indicate *in vivo* and explant respectively. Numbers next to the letters indicate the somite number as shown in (b). Somite 1 is the last formed somite. Scale bar, 100 $\mu$m.

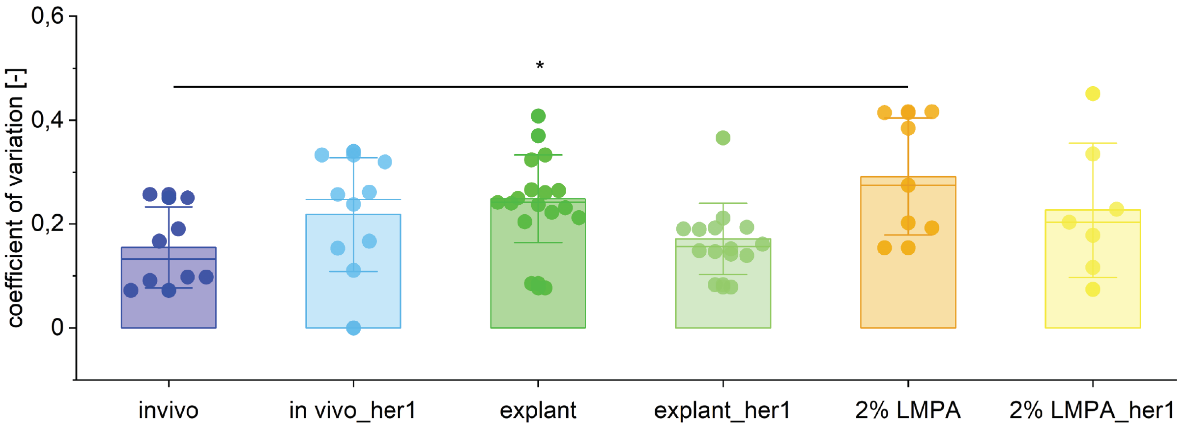


**Supplementary Fig. 5 Coefficient of variations of periods reported in Fig. 2b and c.** Coefficient of variation for individual samples of *in vivo*, free explant and embedded explant. In somitogenesis period *in vivo* samples have lower coefficient of variation compared to free and embedded explants while for segmentation clock period (name_her1) coefficient of variations of *in vivo*, free and embedded explants are similar. Also, variation between somitogenesis and segmentation periods are not significant.

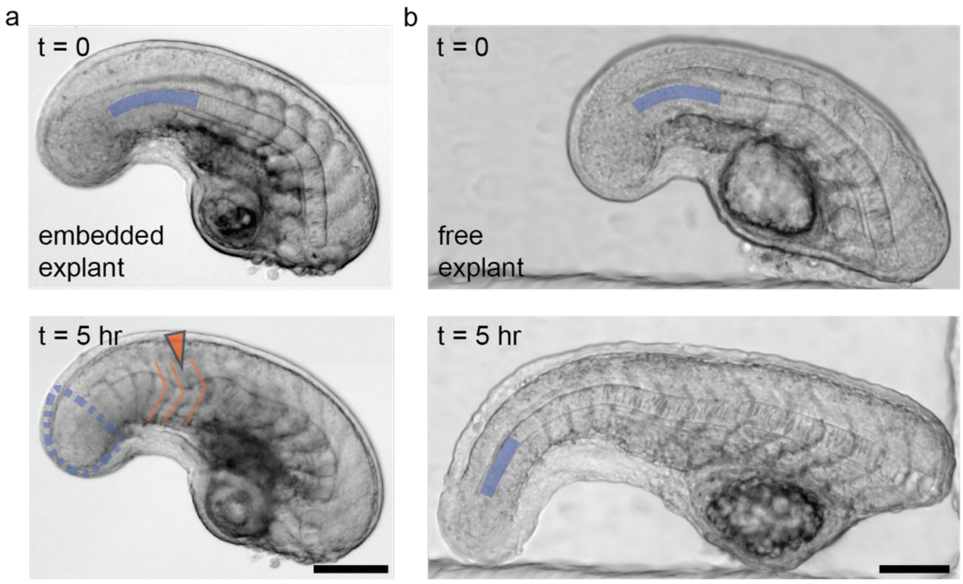


**Supplementary Fig. 6 Embedded explants have reduced PSM length due to prohibited elongation yet they segment and segments take the chevron shape. a** Embedded explant, **b** free explant at t = 0 and 5 hours. Blue lines show the length of the PSM which is significantly reduced from t = 0 to 5 hours, in

(a) compared to (b). In (a) at t = 5 hours, unsegmented region is highlighted with the dashed blue contour however, it is not clear the distinction of the PSM and tailbud. Orange lines show the chevron shape of the somites formed after embedding. Orange arrow indicates the location of buckling. Scale bars, 100 μm.

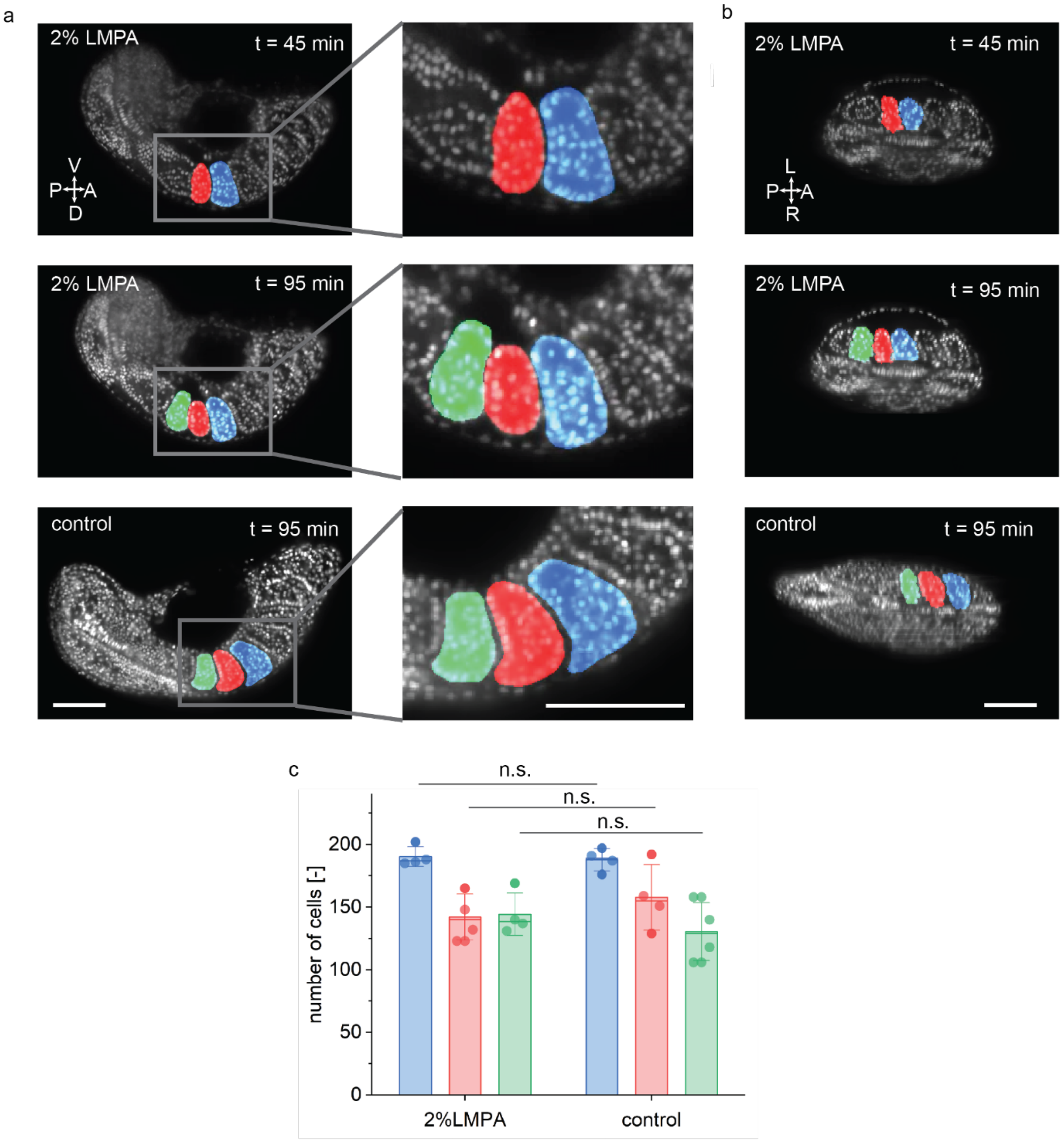


**Supplementary Fig. 7 Comparison of number of cells in somites and shape of somites for explants freely grown in medium and explants embedded in 2% LMPA. a** Example of a H2B-mCherry expressing explant growing in 2% LMPA at 45 minute (top), and at 95 minutes (middle) after the embedding. An example free explant growing in medium at 95 minutes which corresponds to the stage of the embedded explant (bottom). **b** The samples in (a) rotated 90° around AP axis. Corresponding somites are highlighted with the same color. Scale bars, 100 μm in all panels. **c** Number of cells counted in each somite right after their formation for 2% LMPA embedded (n = 4) and free explants (n = 4). Colors of the bars correspond to the highlighted somites in (a) and (b).

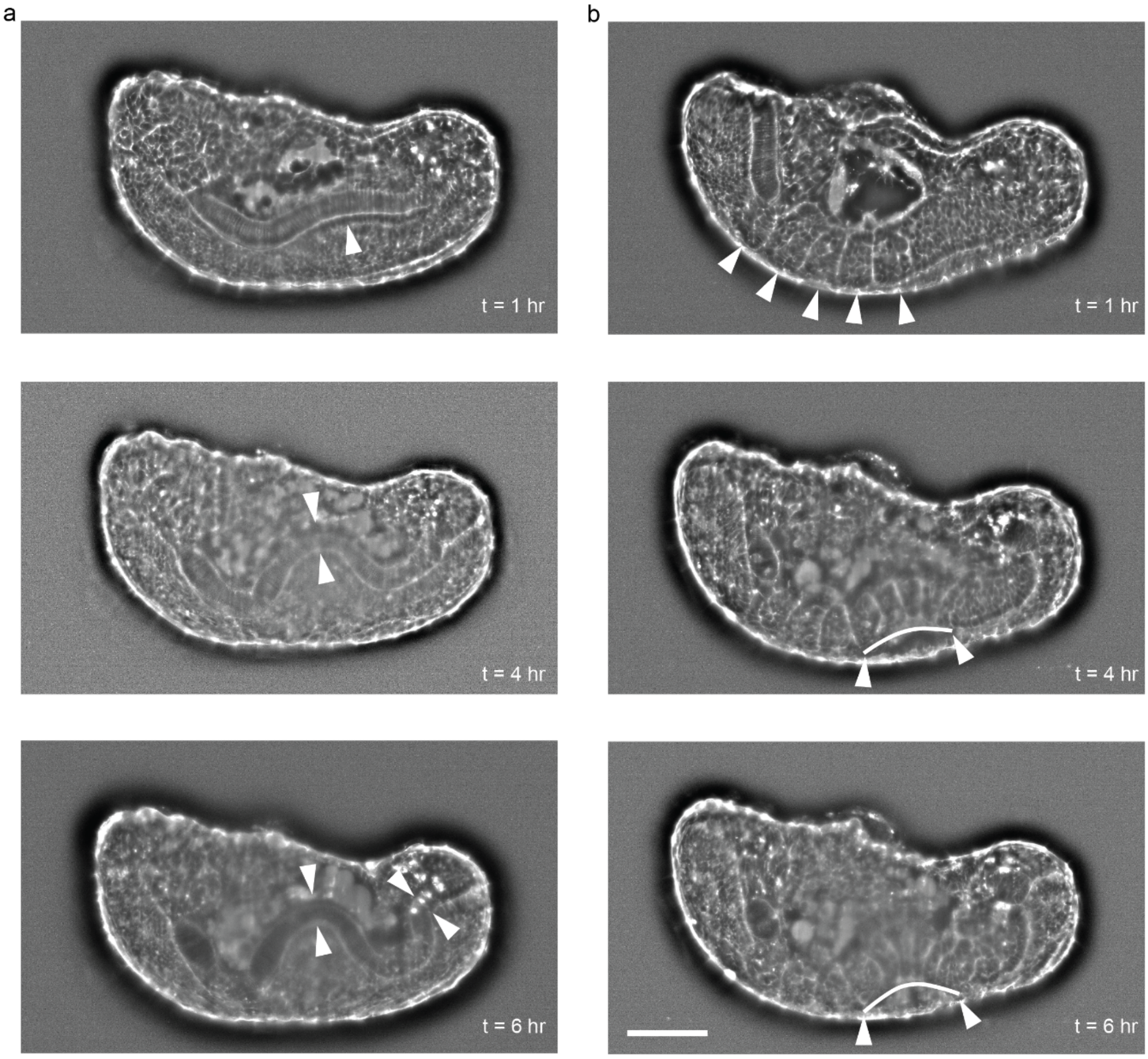


**Supplementary Fig. 8 Shape change of the notochord and somites in embedded tail explant of an embryo from utr-GFP transgenic line. a** Shape change of the notochord. Increasing bending can be followed by looking at the white arrow heads. In the bottom image (t = 7 hr) starting of a second buckling can be observed. **b** Shape of somites. White arrow heads and lines indicate the somites. Top image shows the somites that were on the explant after embedding together with the ones newly formed during the 1st hour of embedding. Middle and bottom images show the abnormal shape of the somites forming while the tail is embeded to 2% LMPA. Scale bar, 100 $\mu$m.

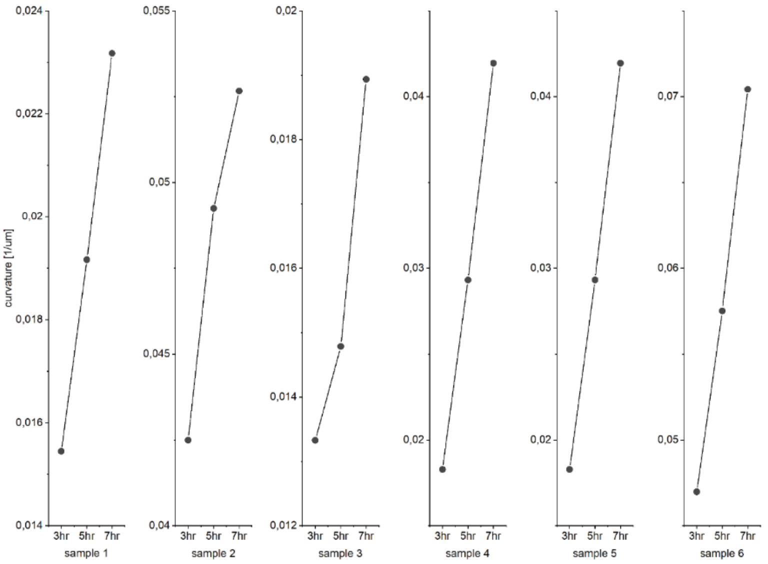


**Supplementary Fig. 9 Curvature increase showing the increase in the degree of bending at the site of buckling over time in the agarose embedded tail explants (n = 6).**

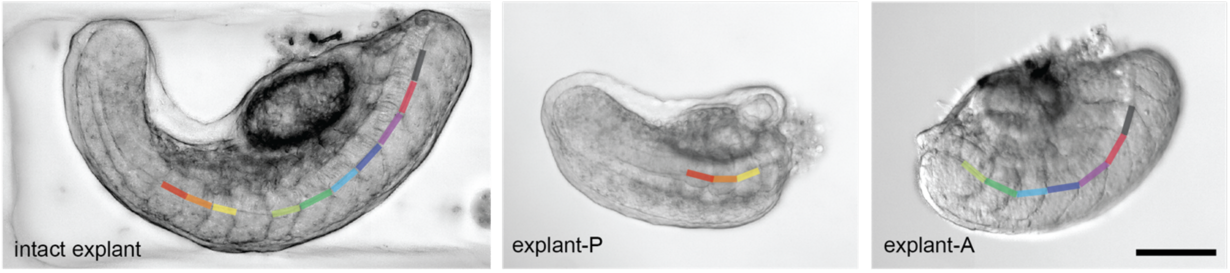


**Supplementary Fig. 10 Somite sizes are similar in intact tail explant and explant-A and explant-P.** Coloured lines with identical lengths show the corresponding somites in each sample. Scale bar, 100 µm.

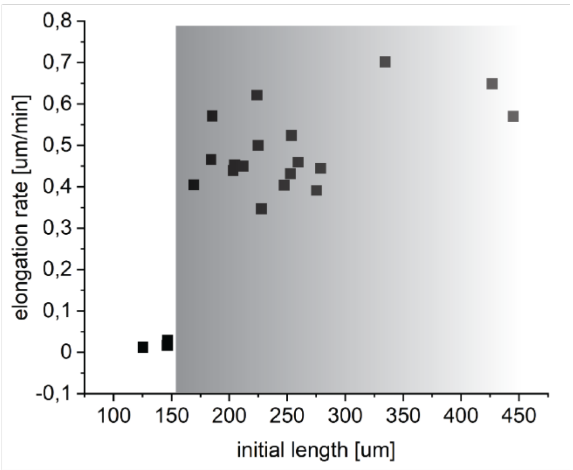


**Supplementary Fig. 11 Initial length vs. elongation rate of explant-Ps.** Explant-Ps that are shorter than 150 µm do not elongate (n = 4), while above this size, they elongate with similar rates (n = 17).

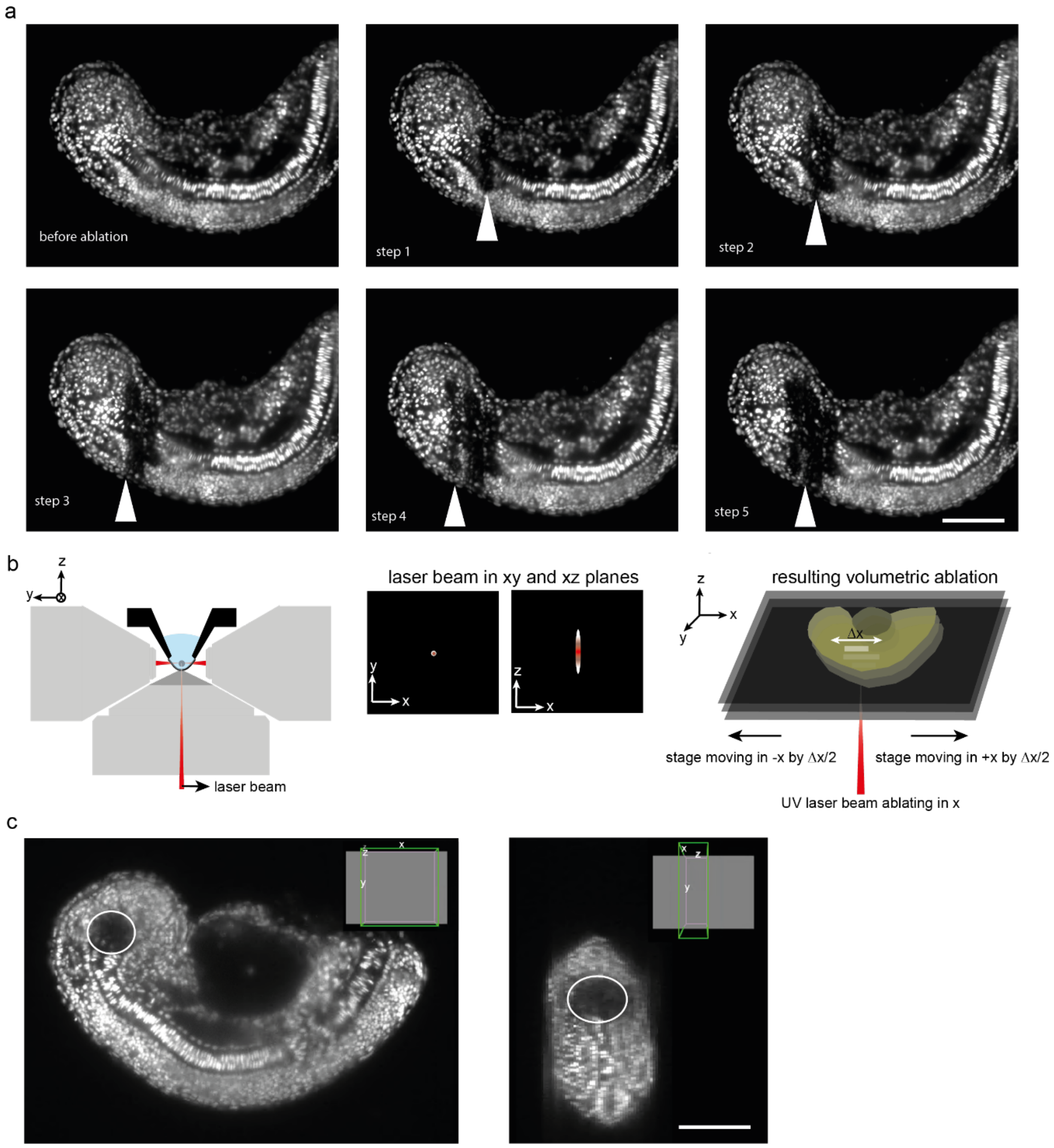


**Supplementary Fig. 12 Laser ablation process. a** An example laser ablation test-experiment showing 5 successive steps involving moving the location of the ablating beam. Each step shows one sweep of the ablation laser across the axis, focused at the plane of the imaging light-sheet. The ablated area is much larger than the one used in the experiment to ablate the progenitor domain, shown in (c). **b** Schematic showing the UV laser's ablation direction on the light-sheet microscope to describe the point spread of the laser beam in different planes resulting in 3D ablation (not to scale). **c** Progenitor domain ablation experiment illustrating the way the point spread function of the laser beam results in a localized 3D volumetric ablation. Scale bars, 100 μm in all panels..

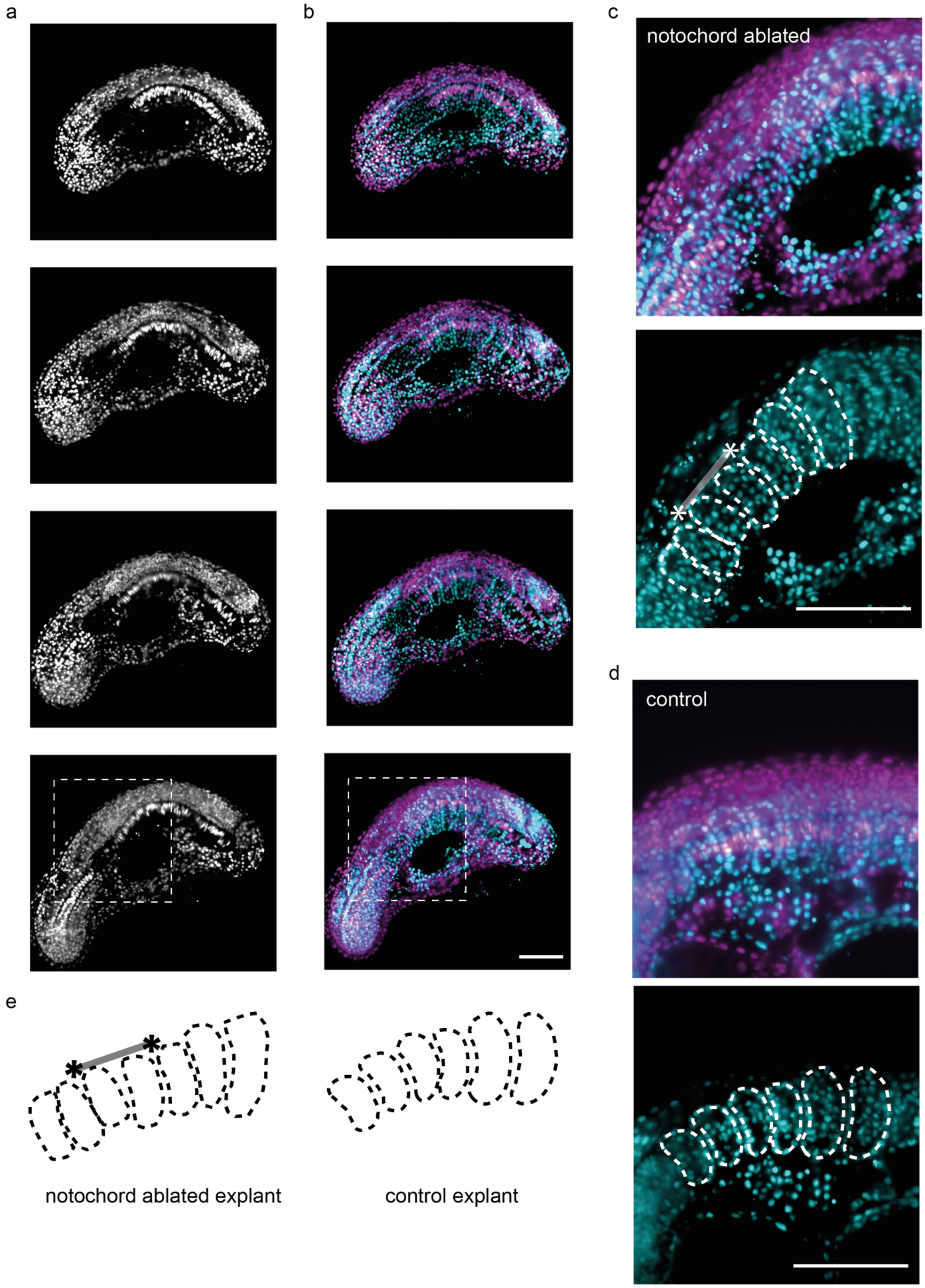
a
b
c
notochord ablated
d
control
e
notochord ablated explant
control explant

**Supplementary Fig. 13 Somite shape in the notochord ablated explants. a** Image sequence focusing on NC. **b** Merge of images focusing on NC (magenta) and somites (cyan). **c** (top) Zoomed in version of the last image in (b) indicated with white rectangle. (bottom) Somite boundaries are indicated with the dashed lines and ablated region of the notochord is indicated with the line between two asterisks (*). **d** (top) A control explant from the same region and same developmental stage as the notochord ablated explant in (c). (bottom) Somite boundaries of the explants are highlighted with dashed white lines. **e** Somite boundaries of the samples in (c) and (d). Scale bars, 100 μm in all panels.

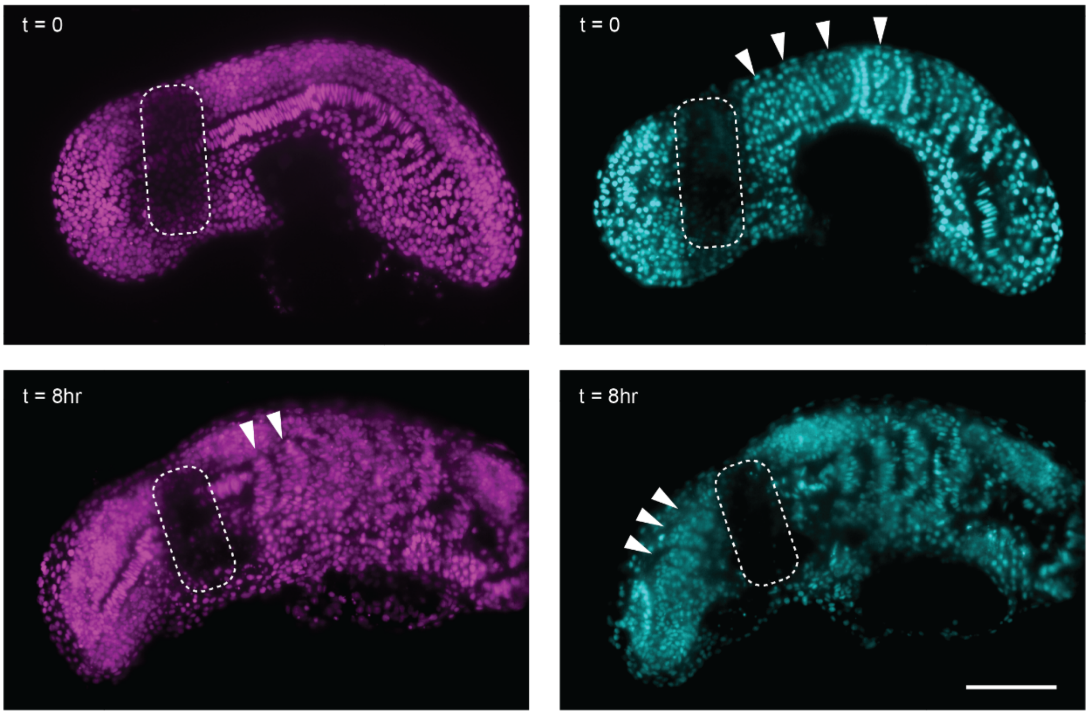


**Supplementary Fig. 14 Bilateral PSM ablation.** Image sequence of both sides of the PSM ablated tail explant. Magenta images are at the focal plane of the notochord showing the elongation of the notochord and cyan images are at the focal plane of the somites showing the shape of the existing and newly formed somites (white arrows). Dashed white contour indicates the ablated region. Scale bar, 100 μm.

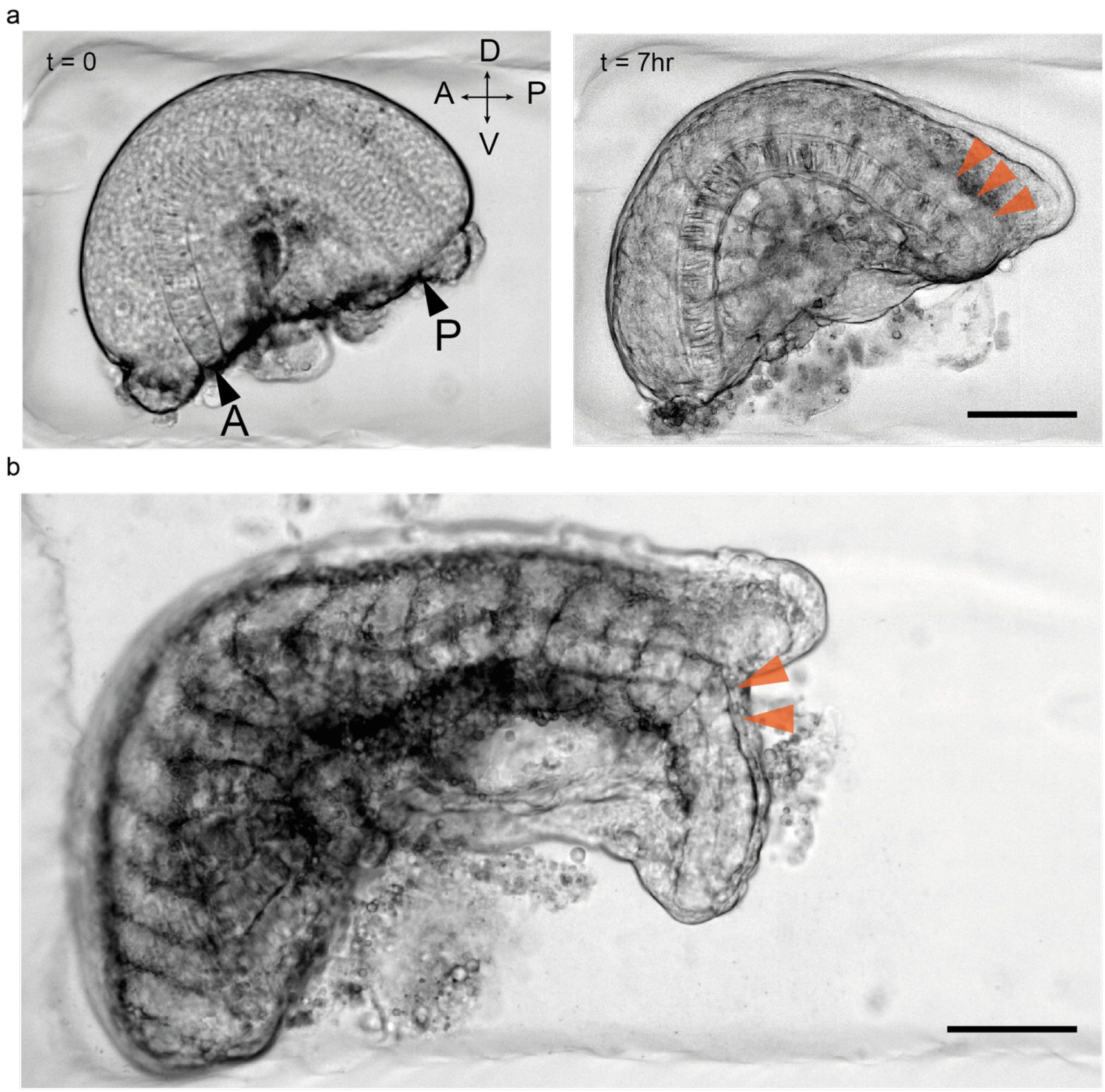


**Supplementary Fig. 15 Somite formation after the micromanipulation of the tailbud of the tail explants. a** Remaining PSM is segmented fully in the explant piece without the tailbud. Black triangles indicate A: anterior and P: posterior ends of the explant. Orange triangles at t = 7 hours indicate the three last formed somites. **b** Dorsoventrally smaller somite formation in the bifurcated tail. Orange triangles indicate the most recently formed two somites in the bifurcated part of the tail to show dorsoventrally smaller somites. Scale bars, 100 μm in all panels.

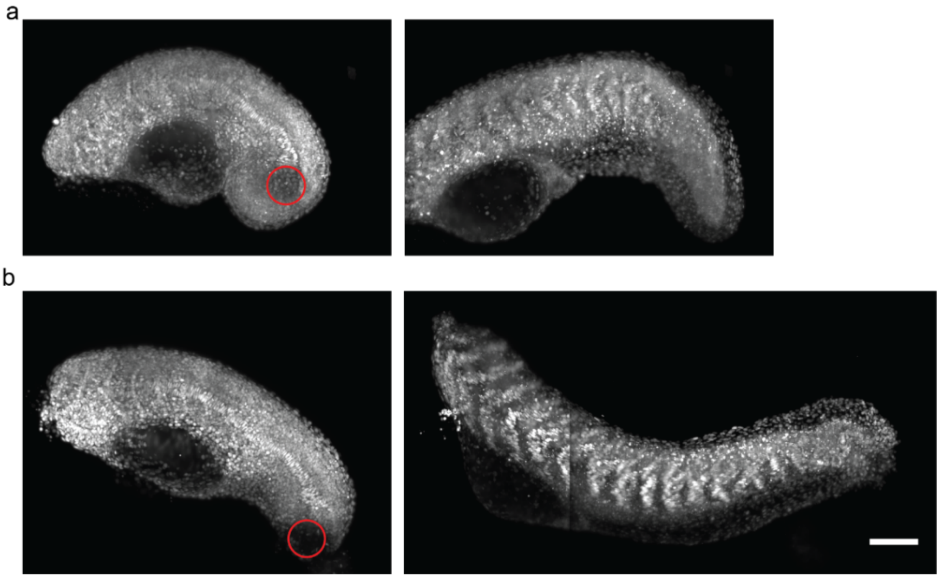


**Supplementary Fig. 16 Notochord progenitor domain ablated samples have shorter final length than tip of the tailbud ablated samples.** Red circles indicate the ablated region. Right images are taken 10 to 12 hours after the ablation (after the completion of somitogenesis). Max projection images of (a) the notochord progenitor domain and (b) tip of the tailbud ablated samples. Scale bar, 100 μm.

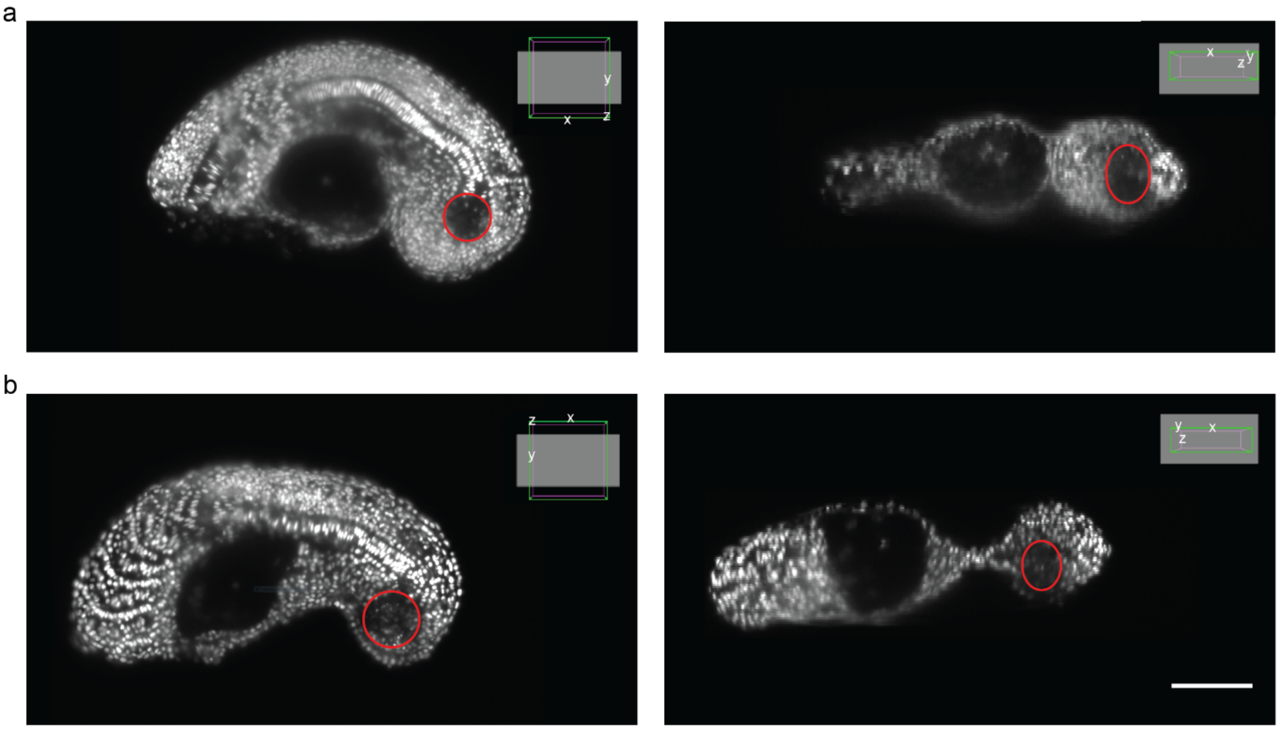


**Supplementary Fig. 17 Volume of the ablated regions are kept close to eliminate the effect of ablated region's size. a** Notochord progenitor domain ablated sample's xy-plane view (left) and xz-plane view (right). **b** Tip of the tailbud ablated sample's xy-plane view(left) and xz-plane view (right). Red circles indicate the ablated region. Scale bar, 100 μm.

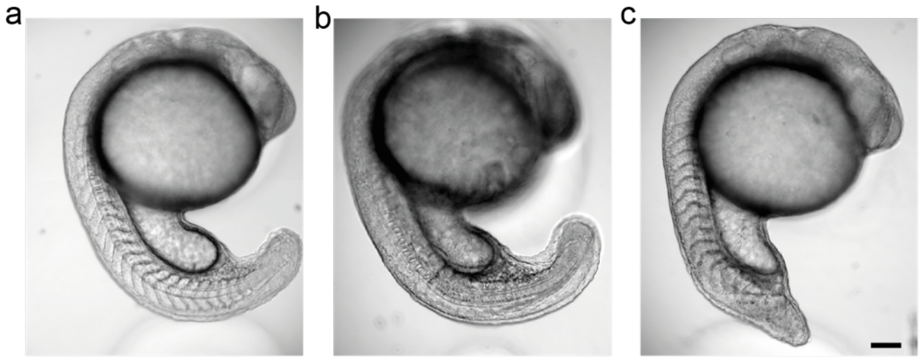

**Supplementary Fig. 18 Control and mutant embryos at the same developmental stage. a** Wild type-AB embryo as a control. **b** *her1*$^{+/-}$*her7*$^{+/-}$ mutant embryo. **c** *no tail* mutant embryo. Scale bar, 100 $\mu$m.

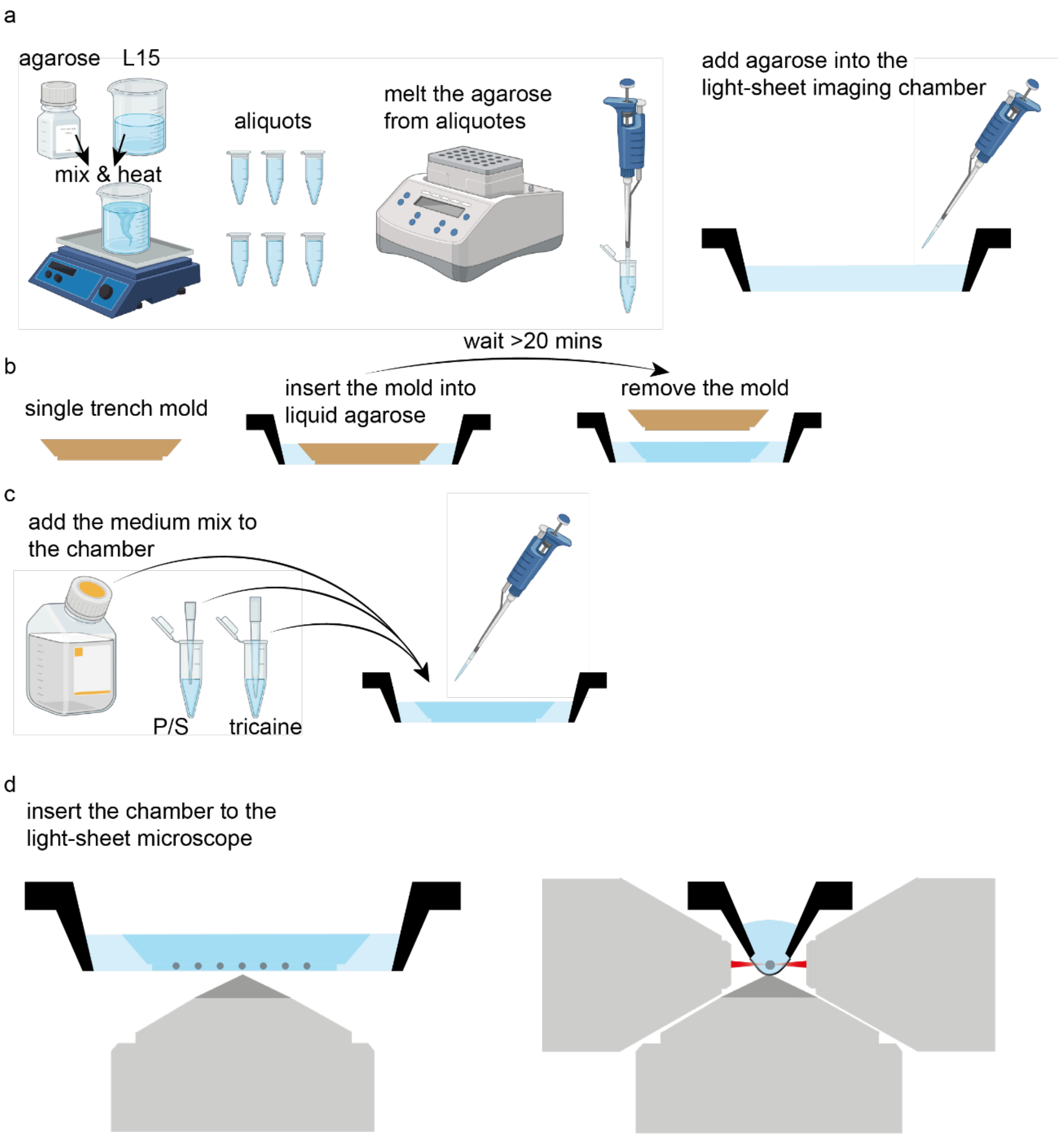


**Supplementary Fig. 19 Chamber preparation process for the light-sheet microscopy Viventis LS1 Live. a** Preparation of the agarose gel. **b** Preparation of the imaging chamber. **c** Imaging medium components. **d** Schematic side views of the chamber inserted into LS1 together with objective and 2 light sheets.

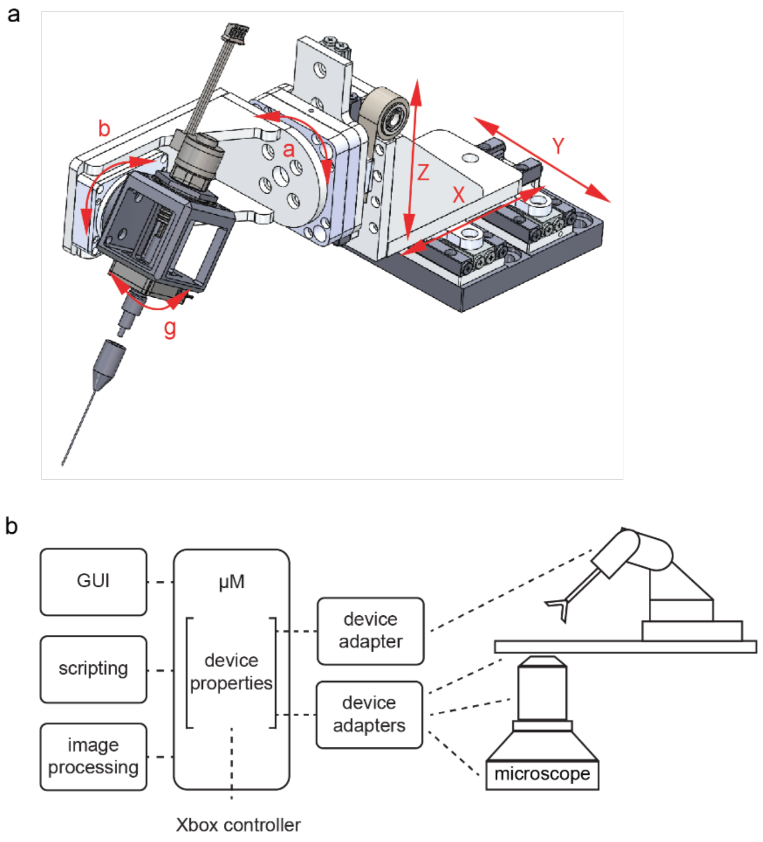


**Supplementary Fig. 20 Robotic surgery of the embryo. a** 3D drawing of 6-DoF robotic micromanipulator with its movement axes. **b** Complete box description of the microsurgery setup.

### Supplementary Movies

**Supplementary Movie 1: Zebrafish embryo tail explant growth.** Bright-field time-lapse movie of a tail explant extending and elongating for 5 hours.

**Supplementary Movie 2: Twitching of tail explant.** Bright-field movie of a twitching tail explant which completed somitogenesis.

**Supplementary Movie 3: Buckling of the notochord of an embedded tail explant.** Example of a H2B-mCherry expressing explant growing in 2% LMPA indicating the buckled location's anterior progression every 1 hour for 8 hours.

**Supplementary Movie 4: Balling up of a small posterior explant.** Bright-field time-lapse movie of a tail explant further dissected into anterior and posterior parts.

**Supplementary Movie 5: Bilateral PSM ablation.** Elongation of the tail and the notochord of a H2B-mCherry expressing explant for 8 hours. Both sides of the posterior PSM are ablated.

**Supplementary Movie 6: Her1-YFP signal of an explant after bilateral PSM ablation.** The expression of Her1-YFP signal in a tail explant. Signal is lost when the wave front passes the ablated region.

**Supplementary Movie 7: Robot assisted microsurgery.** A live demonstration of cutting tail explants from zebrafish embryos using the robotic manipulation platform.

### Supplementary Note. Robotic Micromanipulation

**Precision and accuracy.** The translation stages are linear stages with a parallel-rail structure. One rail is the stick-slip actuator and the other a passive guide. Z has in addition a constant-force spring to compensate for the weight of the spherical wrist. α and β are closed-loop rotary stages and γ is an open-loop rotary stage. The linear stages have a range of 40 mm and closed-loop resolution of 4 nm. They can be back-driven by applying a force of 5 N and apply maximum forces of around 4 N during motion. The rotary stages α and β have a resolution of 25 μ° (microdegree), can be back-driven by applying torques of, respectively, 15N-cm and 6 N-cm and apply maximum torque of, respectively, 6 N-cm and 3 N-cm. γ has a resolution of 3 m° (millidegree). The maximum stage speed is around 20 mm/s for the translational stages, 15 degree/s for α and β and around 45 degree/s for γ. The stepper motor has a per step increment of 0.02 mm delivering up to 35 N of force.

**Referencing**. The zero positions of the translation stages are set by their mechanical end stop. The α and β stages are equipped with a sensor that has nanometre resolution and a reference mark. While α has mechanical stops to prevent full revolution, safeguarding rotation in β required a specific software implementation. To ensure collision free referencing, the range is restricted to β = [0°, 90°] with a defined shutdown position at 0°. The open-loop rotary actuator γ does not have a mechanical end stop for operational reasons and is currently excluded from the referencing operation.

**Software.** The Smaract MCS controller unit handles the host communication and controls all stages. A μManager hub object is used as a single communication port to the controller to implement all cross-channel logic (Supplementary Fig. 20b). The linear stage class of μManager is used for all six stages due to the lack of a rotary stage class. Although the Arduino microcontroller has a simple serial connection protocol that is used by a generic device adapter class, it is accessed by the hub object to retrieve stepper motor positions. Device adapters communicate with μManager by exposing a number of properties which μManager can, respectively, read or write and potentially trigger an action of the connected device. A property is a field consisting of a name and a value. Properties can be accessed within the built-in GUI, via the beanshell scripting environment, or implemented plugins. We developed a user-friendly μManager-based GUI Plugin for manual command entry and retrieval of recorded data. The overall framework is extendable to closed-loop control, particularly based on visual feedback thanks to the availability of a rich computer vision library.

**Motion control**. Teleoperation is performed using an Xbox controller connected to the µManager with the ASI Gamepad plugin (https://micro-manager.org/ASI_Gamepad_Plugin). Xbox controllers are optimised for multiple DOF motion control. Button assignment is customizable, depending on the specifics of the manipulation task. There are two joysticks and we assigned them to control either the translational stages or the spherical wrists. Upon displacement of the joystick, a discrete property value between -1 (left or down) and 1 (right or up) is sent to the corresponding stage. With increasing joystick displacement, the corresponding stage moves at around 25% of the maximum speed and linearly increases to 100%. To eliminate the errors accidentally caused by hand shaking, a displacement threshold is set. This way, parasitic movements occurring with less than a given relative movement amplitude are filtered out.

For tool actuation, the stepper motor is controlled using either a dynamic or a discrete mode. Dynamic control follows the same principle as stage movement, whereas an intensity vector that linearly goes from the zero to the maximum position is assigned to a trigger on the Xbox controller. Upon trigger release the stepper motor goes back to the zero position. The discrete method divides the range into steps and a trigger of a button changes a property value that indicates the discrete position of the stepper motor. The zero position, step range, and maximum position of the stepper motor are set and can be changed within the software.

**Speed layer**. Depending on the type of experiment and location of the manipulator, a variety of velocities or sensitivities are required, respectively. Speed layer functionality was implemented as a property value (1, 10, 100, or 1000), which acts as a simple multiplication factor of the speed variable that is computed on each movement based on the joystick displacement.

**Motion recording and replay**. The manipulator is equipped with a teach-in and replay functionality. Recording mode is activated by changing a Boolean property value. On activation, each change of position triggers the printing of all seven positions into a text file. The replay function runs the motion pattern from a selected text file. The motion pattern can be replayed from 0.1 up to 10 times of the original speed and offsets can be added to the translation stages.

For serial sample processing, the program has record and replay functions and is integrated into µManager. µManager provides a large ecosystem of existing device adapters for over 200 hardware drivers. The software platform specifies a device adapter API and unifies the driver interfaces from imaging devices to motorized stages of different vendors. The code is written in C++ programming language and it is amenable to future extensions such as solving the inverse kinematics problem and implementing elaborate referencing schemes. The device adapters are used as building blocks for various experimental units (e.g. microscope stage, syringe pump) on three different levels: (1) Individually compiled device adapters can be linked directly against a C/C++ application, independent from the µManager core functionality. (2) The GUI of µManager is used to design and script (beanshell) experimental protocols including imaging and automated image processing. (3) Using one of the µManager wrappers for MATLAB or Python, the capabilities of specialized software can be tapped on such as machine learning (e.g. Pycro-Manager).
